# Capturing the liquid-crystalline phase transformation: Implications for protein targeting to sterol ester-rich lipid droplets

**DOI:** 10.1101/2022.06.05.494869

**Authors:** R. Jay Braun, Jessica M.J. Swanson

## Abstract

Lipid droplets are essential organelles that store and traffic neutral lipids. The phospholipid monolayer surrounding their neutral lipid core engages with a highly dynamic proteome that changes according to cellular and metabolic conditions. Recent work has demonstrated that when the abundance of sterol esters increases above a critical concentration, such as under conditions of starvation or high LDL exposure, the lipid droplet core can undergo an amorphous to liquid-crystalline phase transformation. Herein we study the consequences of this transformation on the physical properties of lipid droplets that are thought to regulate protein association. Using simulations of different sterol-ester concentrations we have captured the liquid-crystalline phase transformation at the molecular level, highlighting the alignment of sterol esters in alternating orientations to form concentric layers. We demonstrate how ordering in the core permeates into the neutral lipid/phospholipid interface, changing the magnitude and nature of neutral lipid intercalation and inducing ordering in the phospholipid monolayer. Increased phospholipid packing is concomitate with altered surface properties, including smaller area per phospholipid and substantially reduced packing defects. Additionally, the ordering of sterol esters in the core causes less hydration in more ordered regions. We discuss these findings in the context of their expected consequences for preferential protein recruitment to lipid droplets under different metabolic conditions.

## Introduction

Lipid droplets (LDs) are energy storage organelles that contain, traffic, and buffer fluctuations in the availability of neutral lipids, such as triacylglycerols (TG) and sterol esters (SE). Acting as the hubs of lipid metabolism, they not only store and dispense neutral lipids for energy and membrane building, but also harbor metabolic enzymes that participate in lipolysis and lipogenesis.^1–3^ Additionally, LDs can interact with transcription factors and proteins involved in the innate immune system, expanding their significance to other facets of the cellular lifecycle.^4,5^ Malfunctioning of LD metabolic pathways or alteration of LD proteomes can lead to an onslaught of diseases due to lipid imbalances. This includes, but is not limited to, obesity, non-alcoholic fatty liver disease, lipotoxicity, and cancer.^6^

LDs originate in the endoplasmic reticulum (ER) membrane, where primarily TGs accumulate between the bilayer leaflets. At a certain threshold, there are enough neutral lipids to nucleate and form a budding LD that eventually separates from the ER.^7^ The newly formed LD with a diameter of 100 nm – 100 μm can reattach to the ER through a membrane bridge, diffuse through the cytosol and/or interact with other organelles. The LD lifecycle is largely regulated by the proteins embedded in or associated with the phospholipid (PL) monolayer surrounding the neutral lipid core. This monolayer distinguishes LDs from organelles bound by PL bilayers.^2^ LD proteins are grouped into two classes depending on their origin. Class I proteins, recently coined ERTOLD proteins, translocate to LDs from the ER bilayer either during budding or upon reattachment via membrane bridges. They often contain membrane-embedded hairpin-motifs. Class II (CYTOLD) proteins target the LD monolayer from the cytosol typically via amphipathic helices that interact with the LD monolayer.^2^ Some of these proteins can bind to either LD monolayers or ER bilayers, as both membranes generally share the same PL composition. However, proteins can preferentially target one or the other contingent on the metabolic state of the cell and the lifecycle stage of the LD, such as an expanding LD under growth conditions.^2,7^

Identifying how proteins dynamically target LDs to regulate lipid trafficking and metabolism is an emerging area of research. It is somewhat challenging as LDs do not utilize the dedicated protein-targeting machinery or biochemical landmarks used by other organelles, such as those found in peroxisomes, the ER, or mitochondria.^8,9^ Various studies have proposed that targeting mechanisms instead rely on the organelle’s unique physical properties.^10–14^ Early molecular dynamics (MD) simulations suggested that a substantial amount of interdigitation between the TGs and the PL monolayer led to increased solvent exposure of hydrophobic groups called packing defects.^15^ Subsequent all-atom MD simulations spanning 8 μs revealed that certain LD properties require more than 1 μs to reach equilibration.^14^ With sufficient sampling, 5-8% of the monolayer became composed of TG molecules that had intercalated into and aligned with the PLs. These surface-oriented TGs (SURF-TGs) not only act as chemically unique membrane components, but they introduce chemically-distinct packing defects with an exposed glycerol group in place of a PL head group. SURF-TGs, along with the substantial level of interdigitation between neutral lipid and PL tails, significantly increases the size and lifetime of observed packing defects in a LD monolayer compared to a bilayer of the same composition.

Class II targeting was linked to physical properties by Prevost and co-workers who conducted MD simulations and *in vitro* experiments on several amphipathic helices. They found that multiple large hydrophobic groups were required for LD targeting, and that these groups interact with the larger, more persistent packing defects found in LDs monolayers.^11^ Following MD work from Kim and co-workers explored growing/budding LDs, which are known to preferentially recruit specific proteins such as CCTα. With an applied surface tension of 15mN/m per leaflet (representative of a growing LD), the number of SURF-TGs increased to ∼18% of the monolayer composition, revealing an interesting interplay between SURF-TG molecules and surface tension. As the monolayer is stretched, more TG molecules intercalate with the PLs, decreasing surface tension. However, as worse surfactants there is a limit to how much SURF-TGs can decrease the surface tension. Thus, for every magnitude of applied surface tension there is a corresponding average SURF-TG composition. The area per lipid (APL) for the PLs in the LD monolayer also increased by 90.8% due to this surface tension (compared to only 16.4% for a bilayer of the same composition), substantially increasing the number and size of packing defects. It was revealed that the increased packing defects upon expansion, whether in the monolayer or bilayer, enabled preferential targeting of CCTα. Interestingly, the SURF-TG glycerol’s also provided a preferred protein recruitment pathway via hydrogen bonding with a conserved tryptophan.^13^

Class I targeting was also linked to physical properties by Olarte and coworkers who used MD simulations and experimental work to study the ERTOLD protein GPAT4. They found a thermodynamic driving force for targeting by sequence-specific residues that repositioned from unstable regions of the PL bilayer to more stable regions of the LD monolayer, with some forming hydrogen bonds with TG glycerol head groups and/or select water molecules just below the monolayer surface.^12^ Although this collection of work provides a general understanding of how LD physical properties enable class I and II protein targeting, these studies have focused on TG-rich LDs. The impact of the neutral lipid core composition remains largely unknown.

Despite carrying numerous types of neutral lipids, LDs are typically categorized as either TG-or SE-dominant, depending on the cell type or the metabolic conditions.^16–19^ In many cases, cells harbor LDs that are TG-rich, as TGs are a primary metabolic substrate.^20^ However, some cell types contain SE-dominant LDs. For example, macrophage/foam cells contain a sterol-rich environment due to interaction with low density lipoproteins (LDL). Once esterified, those sterols are stored in the LD core.^21^ In adrenal cortex cells, LDs are enriched in SEs which are used as precursors for steroid hormone biosynthesis.^22^ Furthermore, recent experimental work demonstrated that certain conditions such as stress or starvation increase the SE ratios in HeLa, Huh7, and yeast cells.^17–19^ Mahamid and coworkers exposed HeLa cells to stress, through starvation and mitotic arrest, which increased the relative ratio and induced a transition of SEs to a liquid-crystalline smectic phase. A smectic phase is characterized by molecules aligning themselves in an orderly fashion in stacked layers.^17,23^ Through microscopy, they observed multiple concentric rings of smectic SEs below the PL monolayer and surrounding an amorphous phase of the remaining TGs in the middle of the LD core. It was suggested that this phase transition and separation of lipids is directly related to the cellular and metabolic conditions and alters LD dynamics. It was further hypothesized that the increase of SEs was likely due to consumption of TG metabolites, and that the concentric smectic rings alter surface properties in a manner that alters occupancy of certain LD-proteins while restricting the accessibility of lipids in the core.^17^ A subsequent experimental study demonstrated that acute glucose restriction in yeast cells initiated the same SE liquid-crystalline smectic phase transformation in LDs. Proteomic studies additionally confirmed the LD proteome is altered when the LD is enriched with SEs.^19,24^ For example, when the SE smectic phase change is induced, Erg6 and the model class I peptide LiveDrop re-localizing to the ER, while TG lipases Tgl4-mNg, Tgl3-mNg, and Tgl1-mNg remained on the LDs. It was suggested that glucose restriction promotes the mobilization of TGs from LDs through lipolysis to provide energy for peroxisomes and the mitochondria, consequently inducing a phase change that remodels the LD proteome.^19^

Additionally, *in vitro* experimental work conducted by Chorlay and Thiam established that LD cores containing different neutral lipid compositions have varying affinities for amphipathic-helix proteins. For example, their model amphipathic helix had a higher affinity for LDs with a TG core, as opposed to a squalene core.^10^ Mejhert and coworkers recently revealed various proteins that preferentially target either SE-or TG-rich LDs.^5^ For example, the transcription factor Max-like protein X (MLX) seems to only target TG-rich LDs, while Adipose triglyceride lipase (ATGL) can localize on either TG or SE-rich LDs. This is curious, as MLX translocates to the nucleus to control the expression of multiple target genes involved in glucose metabolism. Thus, association of MLX with TG-rich LDs, which blocks its translocation to the nucleus, alters glucose metabolism.^5^ This implies that cellular conditions that alter the neutral lipid core composition toward SE-rich LDs can modulate metabolic pathways by altering the LD proteome. This raises the question of how LDs with SE-or TG rich core compositions can recruit specific proteins, while selectively releasing others.

In this work, we conduct all-atom MD simulations of SE-rich model LDs. Our findings reveal how the neutral lipid core composition can alter the LD surface and core properties. We demonstrate how different SE:TG ratios (50:50 vs 90:10) impart different physical properties. Our simulations span a timescale of 8 μs and capture, for the first time, the amorphous to smectic liquid crystalline phase transformation in the 90:10 SE:TG LD at physiological temperature. The liquid crystalline SEs display distinct ordered layers in the smectic phase. Simulations of the 50:50 SE to TG ratio system retain an amorphous core, but nonetheless present altered surface properties compared to pure-TG LDs. Core SEs in both systems are less surface-active, intercalating far less into the PL monolayer than TGs. Moreover, TG molecules intercalate less (forming less SURF-TG) with increasing SE concentration. Interdigitation of the neutral lipids with the PL tail region is also altered, decreasing in depth and changing in character. As TG intercalation is replaced with ordered SE intercalation in the 90:10 LD there is a significant drop in area-per-lipid (APL) and increase in PL tail packing and monolayer order. The reduced APL and number of SURF-TGs results in a monolayer with fewer packing defects exposed to the cytosol. Additionally, the hydration of the LD core decreases compared to that in a TG-rich LD, partially due to increased order. Our findings reveal how an SE-rich core composition affects the dynamic physical properties of LDs. We close with a discussion of the potential implications for how core composition might influence protein targeting and the LD lifecycle.

## Results

### Liquid crystalline smectic phase transformation

#### Ordering of LD core

To characterize the influence of the neutral lipid composition, we created LD models with a 50:50 and 90:10 SE:TG ratio. The SE we implemented was cholesteryl oleate (CHYO), one of the more common SEs found within cells.^25^ Consistent with previous MD studies, we simulated our 90:10 and 50:50 CHYO:TG systems for 8 μs to ensure adequate equilibration of physical properties.^14^ After 3 μs, there was a notable ordering in the 90:10 system that was not present in the 50:50 system. Although the neutral lipids were originally given random orientations in both systems (consistent with the isotropic phase)(Fig. S1), those in the 90:10 system aligned to form two distinct layers with clear translational order that is representative of a liquid crystalline smectic phase (Figs. 1a and S1) (Movie S1). We calculated the orientational order parameter (*S*) along with two and three-dimensional radial distribution functions (RDFs) between the center of mass (COM) of the CHYO molecule sterol groups to quantify ordering. An *S* value of 0 indicates no ordering (isotropic) and a value of 1 indicates perfect ordering (crystalline) (see *Methods*). The 90:10 simulation started at a negative value of -0.18, meaning the initial random placement of the CHYO was slightly orthogonal to the system director. Over 3 μs, however, *S* steadily increased and converged to 0.81 ± 0.02, consistent with an aligned ordering of CHYO molecules along the system director (Fig. 1c).^25,26^ This value is consistent with previous experiments and simulations exploring smectic orientational order parameters, which tend to fall in the range of 0.7 − 0.95.^26,27^

**Figure 1.**
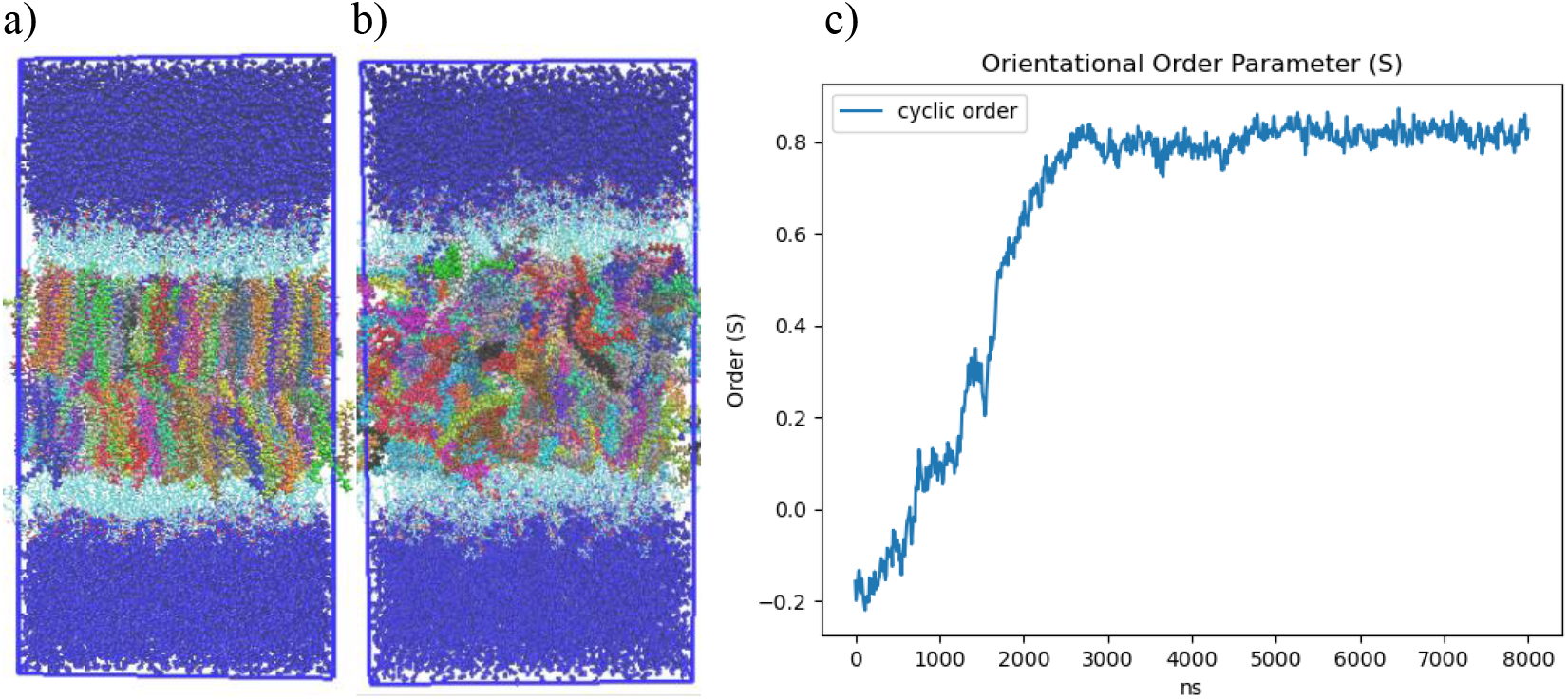
a) 90:10 and b) 50:50 ratio CHYO:TG LD systems after 3 us. The neutral lipids are multicolored to allow for identification of order and disorder c) Nematic order parameter of CHYO in the 90:10 ratio system, converging to a value of 0.81±0.02

Comparing CHYO RDFs from for the first and fifth microseconds of the simulation also demonstrates ordering as a function of the simulation time (Fig. 2). The 3D RDF from the first microsecond has a single peak simply due to co-localization within the LD core (Fig. 2a). That from the fifth microsecond has 3 distinct peaks due to CHYO layering. Similarly, the 2D RDF in the *x*-*y* plane transitions from no peaks (isotropic ordering) to 4 distinct peaks (Fig. 2b). The smectic-phase CHYO molecules align with their oleate tails alternating orientation in an approximate up/down pattern (Fig. S2), as long ago predicted based on crystal structures of sterol esters.^28,29^ This is further supported by approximately equal proportions of “up and down” oriented CHYO molecules per smectic layer (Fig. S2).

**Figure 2.**
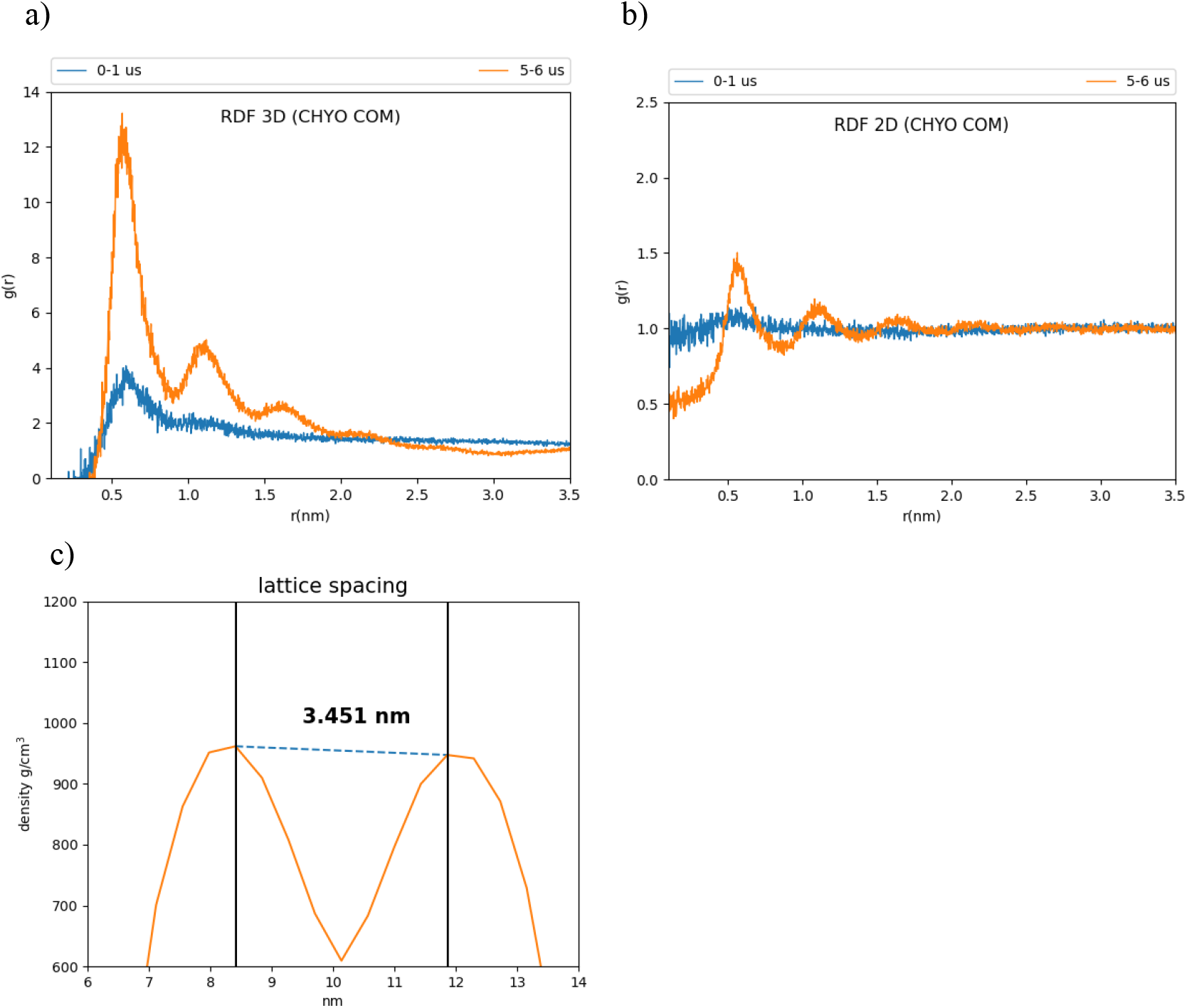
Three and two-dimensional RDFs (a and b) of the 90:10 ratio system. The reference is the CHYO center of mass. (b) The spacing between smectic layers is 3.451±0.035 nm, comparable to experimental results.

In contrast to the 90:10 results, the 50:50 CHYO:TG system retains an orientational order parameter around zero throughout the simulation (Fig. S3). This indicates an amorphous phase with no ordering occurring over the 8μs simulation. Finally, the spacing between CHYO layers in the 90:10 LD system was calculated based on the density profiles of the CHYO smectic phase. The peaks of the profile correspond to the location of the CHYO oxygens (Fig. 2c) due to the overlapping orientation of the tails and side chains (Fig. S2). The spacing of the peaks is determined to be 3.451 ± 0.035 nm (Fig. 2c), which is consistent with experimental measurements of CHYO in LDs (ranging from 3.4 to 3.6 nm).^17–19^

### Physical properties of SE-rich LDs

#### Surface-oriented neutral lipids

SURF-TGs are triacylglycerols that have intercalated into the PL monolayer from the neutral lipid core to behave as chemically unique membrane components (see *Introduction* and Fig S4).^13,14^ Structurally, SURF-TG molecules are similar to PLs with their glycerol moieties oriented toward the water of the cytosol and their acyl tails extending towards the neutral lipid core. Kim and Swanson reported SURF-TGs occupied 5-8% of the PL monolayer of a pure-TG LD simulation.^14^ Our simulations indicate that after an equilibration period of 4 μs, ∼2.5% of the 50:50 system monolayer and less than 1% of the 90:10 system monolayer are composed of SURF-TGs (Fig. 3). CHYO does not appear to be as surface active as TG. CHYO molecules can enter the monolayer for short periods of time but are readily pushed back to the core. They are also relatively few in number, with an average of only 1.3 and 0.21 CHYO molecules (equivalent to 0.96% and 0.15%) existing at the monolayer surface for the 50:50 the 90:10 systems, respectively. Thus, the presence of SE in the LD core reduces the number of SURF-TG proportional to the decreased concentration of TG, and does not intercalate itself to form a surface-oriented SE.

**Figure 3.**
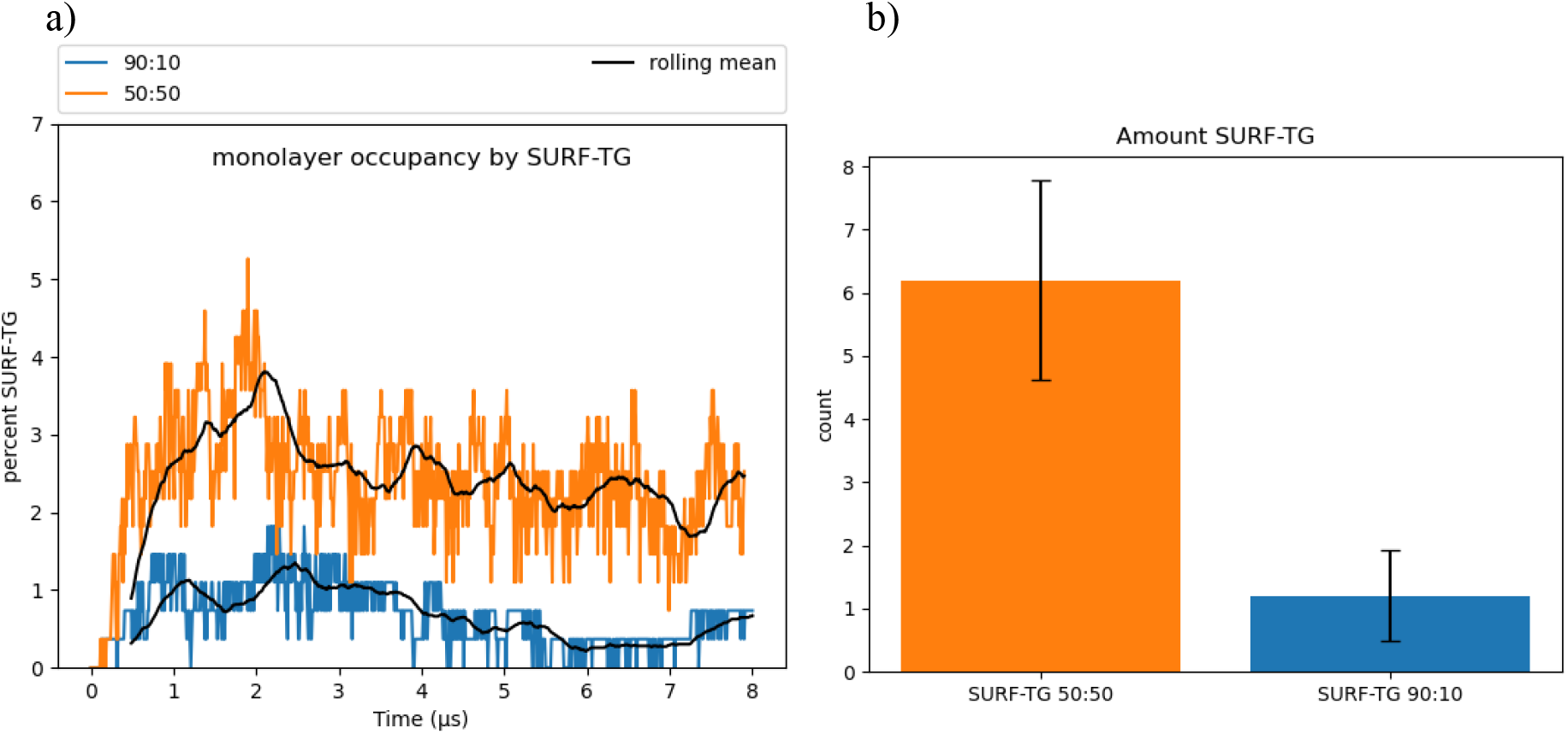
SURF-TGs are not as abundant in SE-rich LDs. Around 2.5% of the 50:50 CHYO:TG ratio system monolayer is composed of SURF-TGs, and less than 1% for the 90:10 ratio system.

#### Interdigitation

The overlap of neutral lipids with PL tails can also alter surface properties. To quantify this for SE-rich LDs, we calculated both the overlap distance (λ_OV_), a traditionally used but approximate quantification of degree of interdigitation, and the relative densities (*ρ*_*rel*_), which more accurately reflect the magnitude of interdigitation (see *Methods*). TG molecules that are found explicitly within the core will herein be called CORE-TG to differentiate them from SURF-TG. Since so few CHYO molecules intercalate into the PL monolayer, we do not distinguish CORE and SURF CHYO. Starting with the 50:50 LD, the density profiles (Fig 4a) show that TG contributes more density than CHYO in the middle of the LD (∼41% CHYO: 59% TG). This is simply a consequence of TG being a larger molecule (885.5 g/mol versus 651.1 g/mol or 42.4%:57.6% of the neutral lipid mass for CHYO:TG). Moving to the region of PL overlap, however, the density profiles from intercalating CHYO and CORE-TG are almost overlapping (Fig 4a) and the total relative density for CHYO increases (∼44%:56% CHYO:CORE-TG). Including the PL density this is ∼19% CHYO: 24% TG: 57% PL (Fig. S5). *This suggests that CHYO molecules intercalate with lipid tails more easily than the CORE-TG molecules, which is perhaps surprising given that TG has three colocalized oleoyl tails to CHYO’s one*. The overlap distance, (λ_OV_ Table S1), which is the integrated overlap parameter *ρ*_*ov*_(*z*) over *z* (Fig 4a bottom), shows equivalent interdigitation distances for CHYO and CORE-TG. Additionally, we find that the overlap parameter profiles for both CORE-TG and SURF-TG are very similar to what was previously reported for pure-TG LDs^14^, suggesting the presence of SEs in an amorphous LD core does not significantly change the nature of TG intercalation into the PL monolayer.

**Figure 4.**
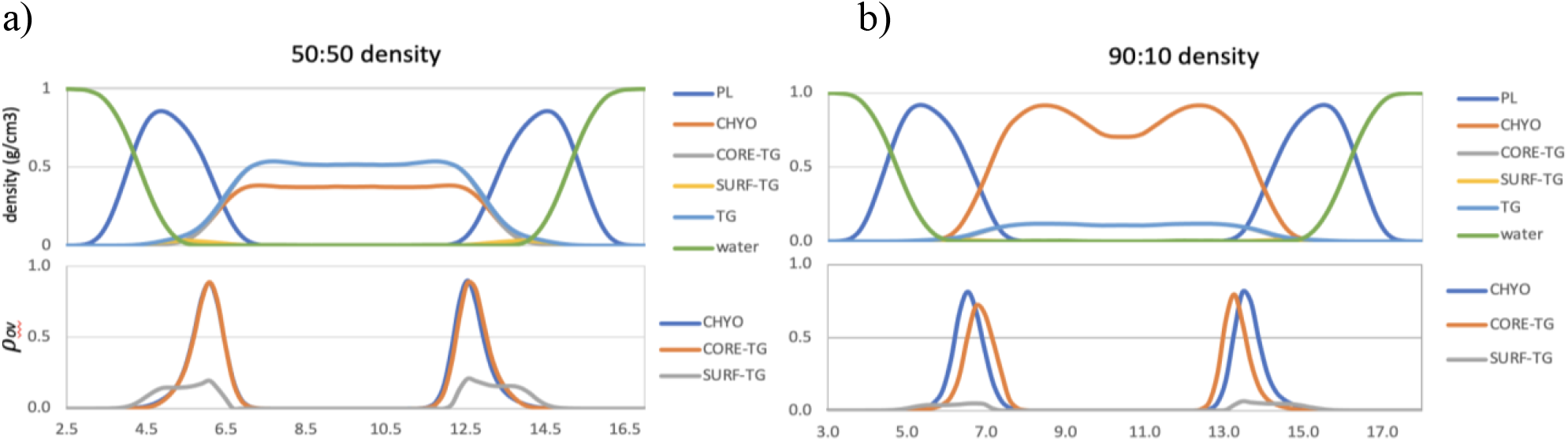
Density profiles (top) and overlap parameters (bottom) for the 50:50 CHYO:TG system (a), and 90:10 CHYO:TG systems (b)

For the 90:10 LD, the overlap distance time-dependence (Fig S6) shows that λ_OV_ of both CHYO and CORE-TG decrease in the 90:10 system as the core undergoes the liquid-crystalline phase transition. Thus, ordering of the liquid smectic CHYO layers influences interdigitation. Once ordered, the density in the middle of the LD (Fig. 4b), which would be 87%:13% for CHYO:TG based on relative masses alone, is now slightly increased in CHYO (∼88%:12%). Consistently, the relative density intercalated into the PL tails is slightly decreased in CHYO (85%:15%), which is equivalently ∼36.5% CHYO: 6.5% TG: 57% PL when including PL density (Fig. S5). Notably, however, the total magnitude of intercalation is not decreased compared to the 50:50 LD. In both systems neutral lipid intercalation makes up ∼42-43% of the density in the overlap region. *Thus, the liquid smectic ordering retains neutral lipid intercalation and pushes some CORE-TG into the PL overlap region and out of the ordered smectic rings*. However, most of the TG molecules are expected to be sequestered into the amorphous LD core under the smectic rings as resolved in microscopy.^17,18^ Interestingly, the overlap parameter also shows that the intercalating CORE-TG in the 90:10 system is shifted slightly down into the LD core and away from the PL-aqueous interface.

Lastly focusing on SURF-TG molecules, they intercalate closer to the PL headgroups with both density and overlap-parameter profiles (Fig. 4) that are consistent with previous findings for pure TG-LDs.^14^ The reduced density magnitudes are expected from the decreased frequency in the 50:50 and 90:10 systems described above. Thus, the phase transition to liquid smectic seems to decrease the number of SURF-TGs, but not to change their alignment in the PL monolayer.

Collectively, for the 50:50 LD we find that: 1) CHYO interdigitates slightly more than CORE-TG but with similar depth, and 2) the presence of 50% CHYO decreases the magnitude of both SURF-TG and CORE-TG intercalation, but does not change their depth profiles when compared to pure-TG LDs. In contrast for the 90:10 LD we can say that ordering of the smectic phase: 1) retains the magnitude of intercalation density, 2) decreases the depth of CHYO intercalation, 3) significantly reduces the number of SURF-TG molecules, and 4) shifts some of the CORE-TG into the PL monolayer (or amorphous core), albeit with a slightly shifted depth profile. The potential biological significance of these changes will be discussed below (see *Discussion*).

#### Phospholipid properties

To characterize the packing and degree of order in the PL monolayer we calculated the area per lipid (APL) and tail order parameters of the PLs. APL is calculated by dividing surface area of a membrane by the number of components. For bilayers the exact relationship between APL and lipid order can be nuanced,^30^ but APL generally decreases with increased lipid packing density, saturated tails and smaller lipid headgroups. The increased packing density of PLs was also previously demonstrated to negatively correlate with the binding affinity of amphipathic helices to artificial LDs,^10^ potentially due to altered packing defects (further discussed below). The APL for the 90:10 system converges to a reduced value of 0.619 nm^2^ after the core gains order (Fig. 5), while the 50:50 system converges to a larger value of 0.67 nm^2^. For reference, the previously reported pure-TG LD had an APL of 0.694 nm^2^, which is significantly larger than that of the ER bilayer (0.628 nm^2^) due to the intercalation of SURF-TG.^14^ There is a positive correlation between the APL and depth of interdigitation (λ_OV_) in the 90:10 system, as indicated by the Pearson correlation coefficient of r = 0.803 (Fig. 5b). As CHYO gains order, its intercalation depth into the PL interface decreases (Fig. S6) and the monolayer packs more efficiently, thereby decreasing the APL (Fig. 5a). In contrast for the 50:50 CHYO:TG system, which does not have any ordering within the core, the reduced APL compared to a pure-TG LD is largely due to the reduced amount of SURF-TG in the monolayer (2.5% vs. 5-8%).^14^ It was previously shown that APL in the pure-TG LD is strongly correlated with the number of SURF-TG. With three oleoyl tails, intercalated SURF-TG molecules take up a substantial amount of room. For the 50:50 system, there remains a correlation between APL and the number of SURF-TG, but it is less significant than it was in a pure-TG LD. This is likely due to the interplay between reduced SURF-TG and the altered intercalation described above.

**Figure 5.**
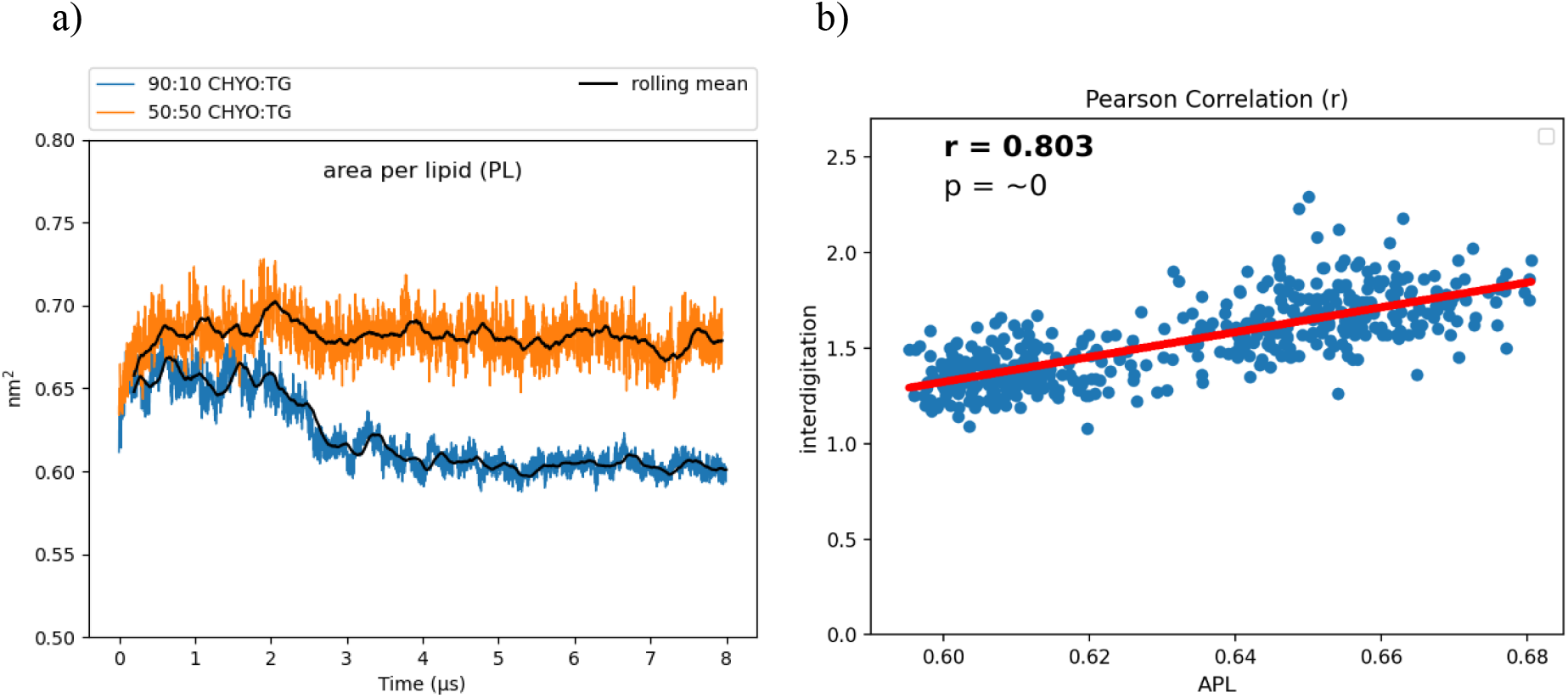
(a) The APL for the 90:10 CHYO:TG ratio LD significantly drops as the system’s core undergoes a phase transition. (b) The APL positively correlates with interdigitation depth, λ_OV_.

Tail order parameters (S_CD_) quantify the degree of order in the phospholipid tails. For POPC and DOPE, the 90:10 CHYO:TG system has significantly increased S_CD_ values compared to the 50:50 CHYO:TG system (Fig. 6) and the ER bilayer (Fig. S7). This increased order in the 90:10 system is expected given the significant decrease in APL combined with an ordered core interface; the PL tails have less area move in as they pack more tightly together. Interestingly, the 50:50 system, which also has a smaller APL due to fewer SURF-TG molecules, has a slight *increase* in fluidity in the PL tail region compared with a pure-TG LD. This is likely due to replacement of approximately half of the CORE-TG intercalation (3 oleoyl tails) with CORE-CHYO intercalation (1 oleoyl tail). Collectively, the APL and order parameters indicate that the 90:10 system has a significantly more ordered and tightly packed PL monolayer than a pure-TG LD, while the 50:50 monolayer has a more moderate reduction in the APL due to loss of SURF-TG and slightly less ordered PL tails due to roughly half of the CORE-TG intercalation being replaced by CHYO intercalation.

**Figure 6.**
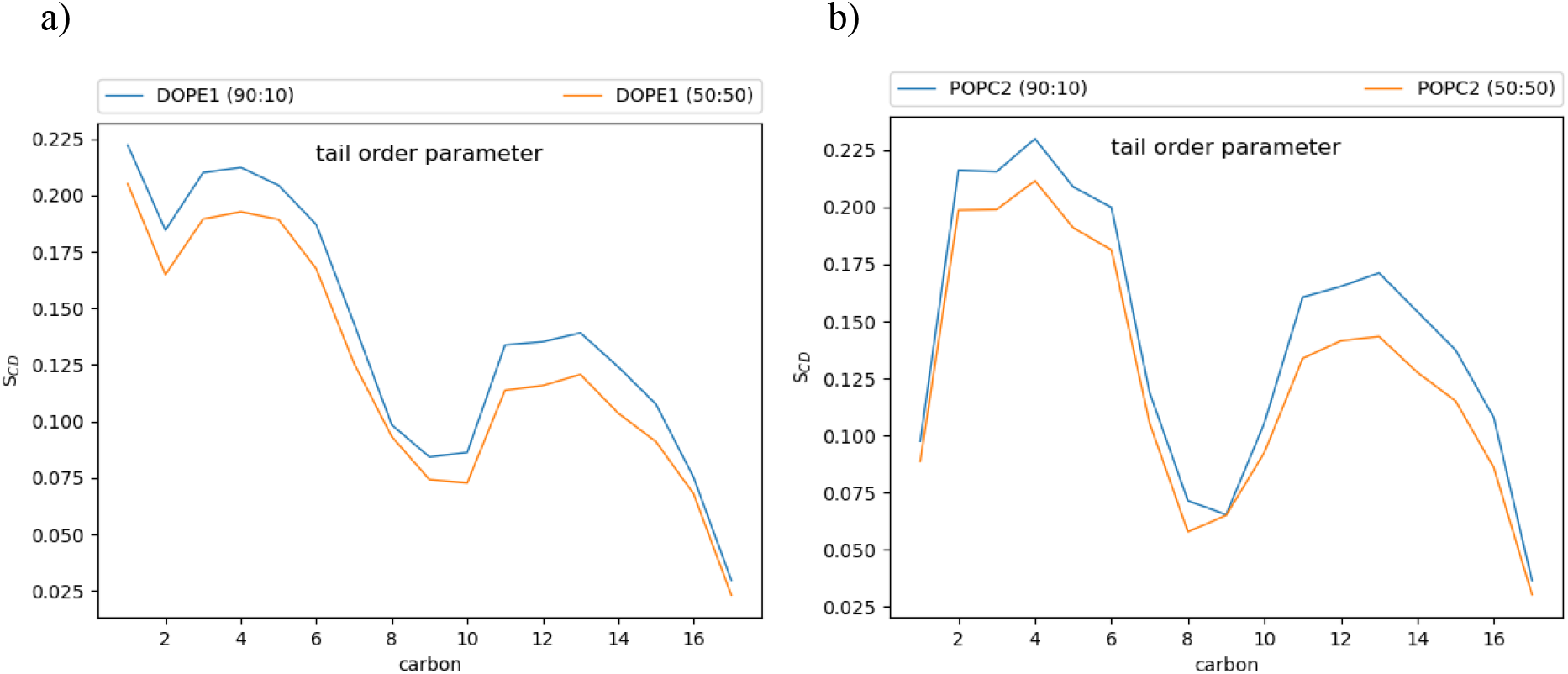
Tail order parameters for DOPE and POPC. In both cases, the 90:10 CHYO:TG ratio LD system is more ordered, due to increased packing concomitate with decreased APL.

#### Packing defects

Packing defects play a crucial role in recruiting amphipathic helices (class II proteins) to both bilayer and monolayer membranes.^2,11,13,31,32^ Bilayers have PL-acyl defects, which are defects that expose the hydrophobic acyl tails of the PLs to the aqueous cytosol. LD monolayers additionally exhibit defects caused by the neutral lipid core. Therefore, we began by distinguishing packing defects based on their molecular composition; PL-acyl, TG-acyl, and CHYO-acyl are defects in which the respective acyl tails are exposed to the cytosol, while TG-glyc are exposed TG-glycerol groups, and CHYO-side are defects caused by either the sterol rings or sidechain of the cholesterol group.

The 50:50 CHYO:TG system has significantly more packing defects as easily visualized when looking down on the monolayers from a bird’s eye view (Fig. 7 a,b). Quantifying this difference with the probability of observing a packing defect as a function of defect size (Fig. 7c) verifies that the 50:50 CHYO:TG system has larger defects. The dominant defect-type for all systems are the PL-acyl defects. To quantify the different types of defects, we calculated packing defect constants (*π*) (Fig. 8), which describe the decay rate of the probability of finding a defect with size N fit to a function *P*(*N*) = *ce*^−*N*/*π*^ (see *Methods*). A higher packing defect constant indicates that there is a higher probability of finding larger defects. In the 50:50 CHYO:TG system the TG-glycerol and TG-acyl defects have slightly higher defect constants than CHYO (Fig. 8c), which is sensible given the presence of 2.5% SURF-TG and nearly identical intercalation of CHYO and TG. Surprisingly, the defects from TG and CHYO are nearly identical in the 90:10 LD, despite the fact that CHYO is present in 9-fold excess and makes up 85% of the intercalated neutral lipid mass (Fig. 8c). This indicates that the few remaining SURF-TG (<1%) and intercalated TG molecules contribute much more significantly to the packing defects than CHYO, potentially due to the broader 3-tail structure of TG compared to streamlined 1-tail structure of CHYO.

**Figure 7.**
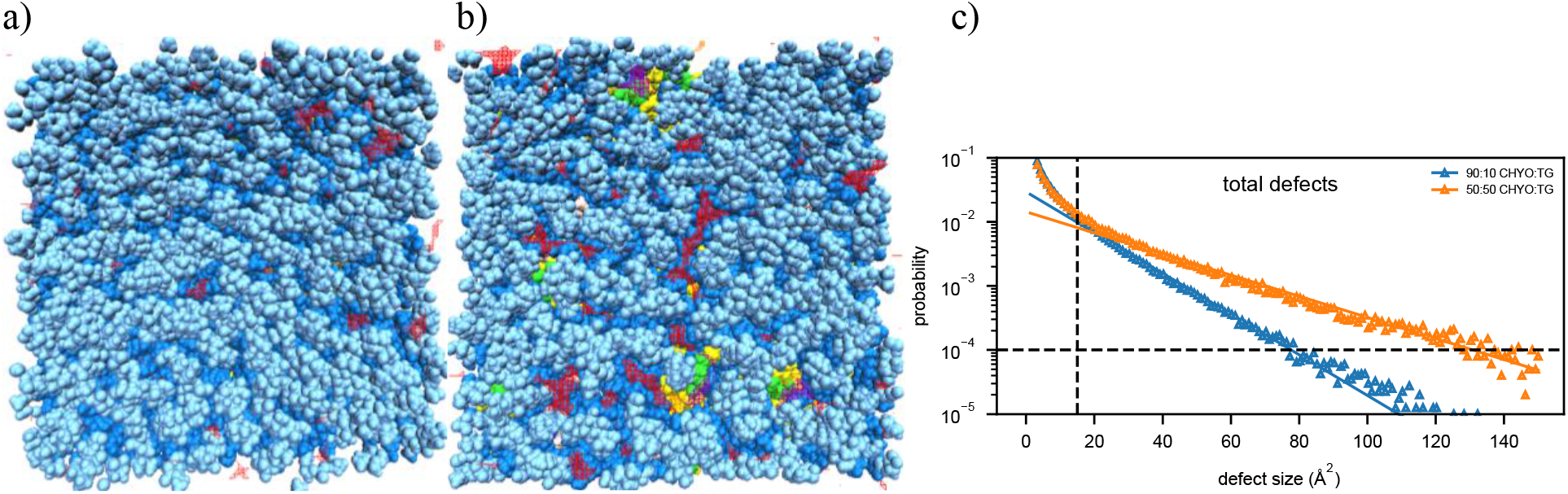
Birds-eye-view of PL monolayer of (a) 90:10 and (b) 50:50 CHYO:TG systems. Light and dark blue are the PL head and acyl groups, while green and yellow are TG glycerol and acyl groups. Red and purple indicate PL-acyl and TG-glycerol defects that are larger than 15 Å^2^, respectively. (c) The probability of finding defects of a given sizes fit to exponential curves highlighting the increased propensity of large defects in the 50:50 LD.

**Figure 8.**
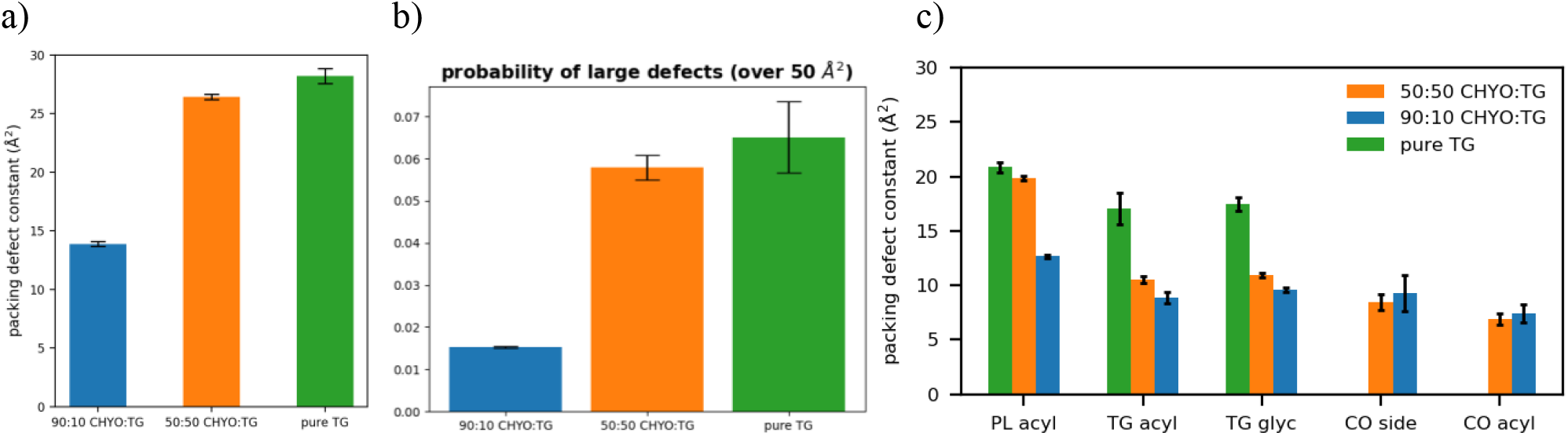
(a) The packing defect constants for the 90:10, 50:50, and a pure TG LDs show that increasing CHYO concentration leads to less defects. (b) The probability of finding a defect over 50 Å^2^ shows the same trend. (c) The respective types of defects demonstrate that PL-acyl are the dominant defects for all 3 systems, while 10% TG induce defects almost as much as the 90% CHYO in the 90:10 ratio LD.

For a broader interpretation, we also calculated the overall probability of any type of defect occurring (Fig. 8a). The 90:10 system has a defect constant of 13.89 ± 0.21 Å^2^ while the 50:50 system has a defect constant of 26.44 ± 0.25 Å^2^. To further compare large packing defects, we then calculated the probability of defects that occur over the size of 50 Å^2^. The 50:50 CHYO:TG system is 3.66 times more likely to exhibit these large defects than the 90:10 system (Fig. 8b). To summarize, the presence of CHYO up to 50% decreases the magnitude of packing defects compared to a pure-TG LD, but not substantially since intercalating CHYO still contribute to packing defects. However, upon transformation to the smectic phase with 90% CHYO, packing defects decrease substantially, with the small relative number of remaining TG molecules contributing more to defects than CHYO. In both cases, PL-acyl defects are the dominant defect type, but even these are reduced with increasing CHYO concentrations.

#### Hydration of the LD core

Our simulations revealed some water molecules penetrate into the neutral lipid core, consistent with previous work on pure-TG LDs.^14^ These results suggest that a higher ratio of CHYO is inversely associated to hydration (Fig. 9), with the 90:10 LD and 50:50 LD systems converging to water densities of 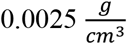 and 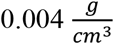, respectively. This is reduced from the previously reported 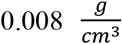 for pure-TG LDs.^14^ It is important to note that these magnitudes of hydration are likely overestimated since the standard CHARMM36 parameters used in our model for TG are not ideal for bulk hydration properties and interfacial (water/lipid) surface tension.^33,34^ The central limitation in these, and all parameters developed to date, is lack of polarizability, which is essential to capture proper neutral lipid interactions in both nonpolar core and polar interfacial environments. For example, bulk TG is hydrophobic, but at a water interface the glycerol moieties should have larger partial charges in response to proximal water. The standard CHARMM36 parameters are on the polar side and thus slightly over-hydrate bulk TG, which was experimentally measured at 1.8 × 10^−3^ g/ml.^35^ However, these parameters correctly capture the APL for the monolayer, which is approximately 15% larger than that of the ER bilayer.^10,14^ This is in large part due to the presence of SURF-TGs, which behave as worse surfactants than PLs consistent with the small surface tension for LD monolayers.^10,36,37^ Updates to the additive forcefields of TG that correct bulk hydration and interfacial water/lipid surface tension do not correctly represent the APL of a LD monolayer, likely due to increased hydrophobicity that retains TG in the LD core.^38,39^

**Figure 9.**
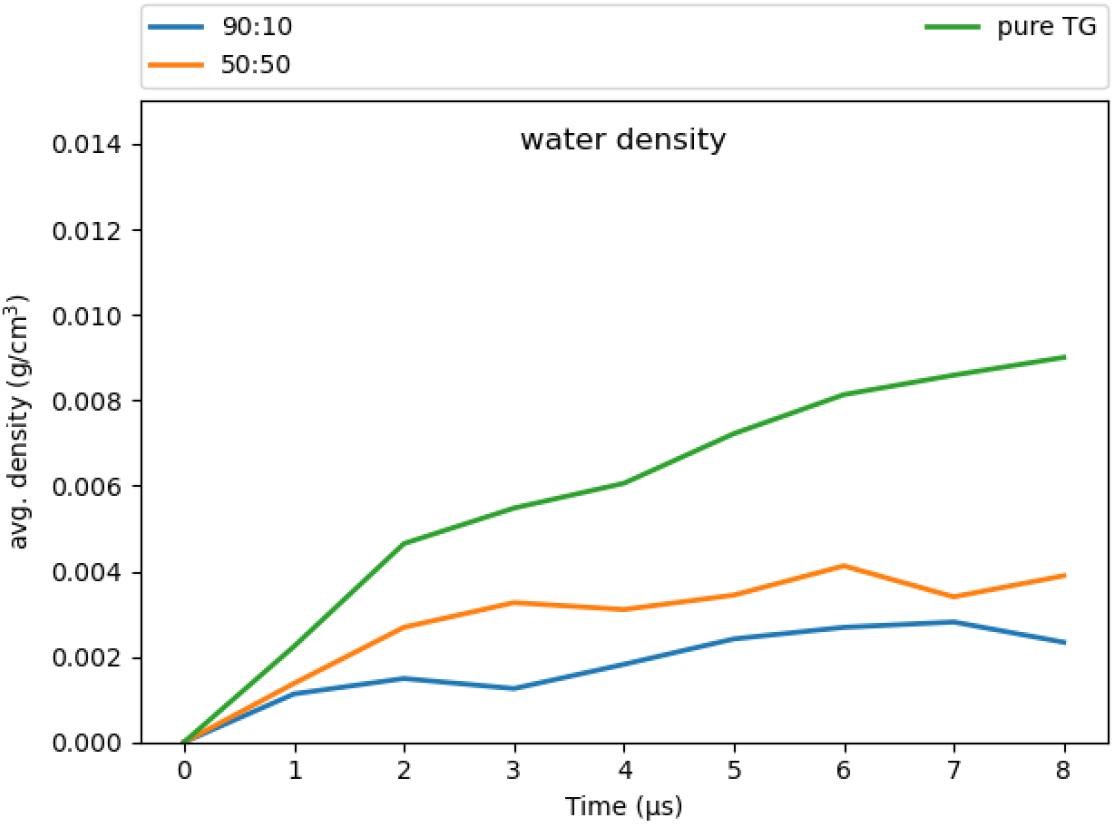
The 90:10 ratio system has the least amount of water in the core, while the pure-TG LD has the most.

The surface properties and APL are in part what make LDs unique organelles, as membrane properties are responsible for recruiting proteins. Since our primary focus is on the LD surface properties, the standard CHARMM36 parameters were selected as most appropriate for this study. Additionally, although the hydration profiles reported herein are slightly overestimated, we anticipate they correctly capture some degree of hydration in the LD core that is enhanced just below the PL monolayer and that decreases with increasing SE/CHYO concentrations. Upon formation of the liquid-crystalline smectic phase any retained core water would localize with the oxygens, pulling back from the PL-neutral lipid interface. These predictions should be compared to experimental analysis of LD core hydration, which would be a very helpful addition to the field, as well as future studies with polarizable force fields.

## Discussion

Recent studies on TG-rich LDs have revealed unique physical properties that are at least partially responsible for protein recruitment. However, there is growing recognition that SE-rich LDs are different and that these differences likely have important implications for metabolic cellular processes and health.^17– 19,21,22^ Notably, it has been shown that when LDs are transitioned to SE-rich cores there is a shift in their proteome as they exclude some proteins while retaining association with others.^5,19,40^ Our objective is to understand how this differential protein recruitment occurs. Using simulations spanning 8 μs, we have herein calculated the physical properties for LDs containing two ratios of SEs to TGs. Our findings provide clues into the mechanisms that would differentiate protein targeting to LDs dependent on their core composition.

Most notably, we observed the anticipated phase change in a 90:10 CHYO:TG LD with the bulk CHYO gaining order over the span of ∼3 μs. The resulting orientational order parameter (S) of 0.81 and lattice spacing between layers of 3.45 nm is indicative of a highly ordered smectic phase.^41,42^ This spacing is also consistent with the 3.4-3.6 nm SE smectic spacing reported from cryogenic electron microscopy.^17–19^ The phase change in our study occurred at physiological temperature, further supporting the recent experimental results suggesting SE-induced phase changes do occur in biological systems.^17–19^ The liquid-crystalline smectic phase within LD cores is distinctive from an amorphous phase of TG-rich LDs, and many of the unique surface properties discussed next stem from this ordering.

It is first important to note that the PL monolayer surrounding a LD of any composition is very different from one leaflet in a PL bilayer. This is partially due to the significant interdigitation of neutral lipids (NLs) into the PL tails complicating the interface. It is also contributed to by surface active TGs (SURF-TGs), which fully intercalate and align with the PLs and act as unique membrane components. We find notable changes in both intercalation and surface-active neutral lipids with increasing SE concentrations.

First, the abundance of surface-active neutral lipids decreases—moderately for the 50:50 LD and significantly the 90:10 LD (Fig. 3). CHYO does not seem to be surface active like TG, so there is no SURF-CHYO equivalent of SURF-TG. This is consistent with the decreased solubility of CHYO in PL membranes compared to TG.^43–45^ Additionally, the number of SURF-TG are reduced from 5-8% in a pure-TG LD to 2.5% in the 50:50 CHYO:TG LD and less than 1% in the 90:10 LD. This will not only impact proteins that directly interact with SURF-TG molecules (discussed with packing defects below), but may also have implications for growth and shrinkage. SURF-TGs act as dynamic membrane components that can modulate surface tension in expanding/budding LDs. Thus, SE-rich LDs may be more resistant to expansion. The CHYO LDs in our simulations are likely comparable to LDs during stressed or starved conditions.^17,19^ Therefore, shrinkage instead of expansion would likely occur, as TGs would be actively consumed as metabolites. The decreased surface tension would in turn decrease the number of SURF-TGs intercalated into the LD monolayer, consistent with our observed decrease in SURF-TGs with increasing SE concentration.

Focusing on intercalation, there is a decrease in the *depth* of intercalation from CORE-TG and CHYO molecules into the PL tails with increasing SE concentration (Fig. S6). Intercalation from CORE-TG is reduced from a λ_OV_ of 1.83 nm for a pure-TG LD^14^ to 1.657 nm for the 50:50 systems, and 1.188 for the 90:10 LD (Table S1). As expected, some of this CORE-TG intercalation is replaced by CHYO intercalation. In fact, the density profiles (Fig. 4) show that the single CHYO tails intercalate slightly more readily than the three colocalized TG tails with nearly identical intercalation depth in the 50:50 LD (Fig. 4a bottom, Table S1). This replacement of TG with CHYO in the PL tail region is a contributing factor to the shift in monolayer packing (APL), order and defects discussed next. For the 90:10 LD, CHYO makes up slightly less than 90% of the intercalation. This is a result of a small number of CORE-TG molecules being pushed into the PL tail region and out of the ordered smectic rings. Similar to CORE-TG, the phase transformation also decreases the depth of CHYO intercalation (Fig. S6, Table S1). However, the total magnitude of intercalation density remains the same as that observed in the 50:50 LD. Thus, the more pronounced changes in physical properties for the 90:10 LD system discussed below are a consequence of the altered depth and shifted composition (mostly now CHYO tails) of neutral lipid intercalation, not due to the exclusion of tail intercalation.

Turning to APL, as the 90:10 LD core gains order, the APL for the PL monolayer also decreases (Fig. 5a). A decrease in APL is generally associated with increased PL packing, but the exact causes can be nuanced. In this case, multiple factors contribute. The first is the decrease in SURF-TG which were previously shown to strongly correlate with the APL in a pure-TG LD.^14^ The drop in APL from 0.694 nm^2^ for a pure-TG LD to 0.67 nm^2^ for the 50:50 LD to 0.619 nm^2^ for the 90:10 LD is consistent with their relative abundance of SURF-TG (5-8%, 2.5% and <1%, respectively). However, reduced depth of neutral lipid interdigitation ((λ_OV_ Table S1)) also contributes, as supported by the correlation between these two variables (Fig. 5b). Lastly, the shift in composition to increased CHYO intercalation combined with reduced conformational flexibility of the smectic phase are expected to be large contributing factors to the decreased APL and attendant increase in PL packing. In the smectic phase, the CHYO molecules are aligned with their tails and side chains oriented in a parallel fashion (Fig. S2), which leads to a more uniform interface with the PL tails. This interface of the interdigitating smectic-CHYO molecules induces increased ordering in the PL tails as indicated by the increased tail order parameters (Fig. 6).

With increasing order and decreasing APL one would anticipate a change in packing defects. This is highly relevant since experiments and MD studies have demonstrated that both APL and packing defects are corelated with the binding affinities of class II amphipathic helix proteins, such as CCT*α* and the amphipathic helix from model protein caveolin 1.^10,13^ Defects enable protein recruitment through exposure of PL-acyl tails and neutral lipids that provide ideal interfaces for class II hydrophobic residues. The small drop in APL for the 50:50 LD, and more significant drop for the 90:10 LD system are consistent with the moderate and significant reduction in packing defects in the 50:50 and 90:10 systems, respectively (Fig. 7,8). Not only are there less defects, but they are also much smaller, with the probability of finding a defect over 50 Å^2^ reduced 3-fold in the 90:10 CHYO:TG system when compared to TG-rich systems (Fig. 8b).

The defect distributions also change. The 50:50 CHYO:TG and pure-TG LDs actually have a similar magnitude of packing defects (26.44±0.25 and 28.62±0.45 Å^2^, respectively), with some of the TG defects of course being replaced by CHYO defects in the 50:50 system (Fig. 8c).^14^ However, TG molecules still contribute more to the packing defects than CHYO molecules in the 50:50 system, likely due to the remaining 2.5% SURF-TG. In the 90:10 smectic LD, the decrease in PL and TG defects is more significant with CHYO contributing far less than 90% of the neutral lipid defects, indicating again that it is less effective in creating defects. These changes in the types of defects present will likely have implications for which proteins are attracted to the LD monolayer. For example, the preferential interactions between glycerol defects and a tryptophan in CCTα, a protein segment that is recruited during LD expansion, could be minimal for a SE-rich LD.^13^ Moreover, the presence of CHYO defects may enable entirely different protein interactions yet to be discovered.

Hydration of the core has previously been proposed to help stabilize certain proteins, such as the class I ERTOLD model peptide LiveDrop.^12^ Specific residues of this membrane-spanning peptide that are emersed in the LD core have charged and polar residues, seemingly unfit for a hydrophobic neutral lipid environment. Coordination with water molecules that are also hydrogen bonded to TG glycerol oxygens could provide the requisite polar interactions to enable such conformations. We find that the degree of hydration for the 50:50 and 90:10 CHYO:TG LDs is decreased compared to that of a pure-TG LD.^14^ This makes sense given that TG has a total of 6 oxygens in its glycerol group, which create a polar assemblage for water interactions. Sterol-esters like CHYO on the other hand, have 2 oxygens, meaning fewer contact points for water. Additionally, the more densely packed monolayer of the 90:10 CHYO:TG LD may make it more difficult for waters to enter past the membrane, which could introduce kinetic barriers for polar protein residues to drag water in. Once inside the core, waters will have less mobility (i.e., stabilizing entropic freedom) due to the ordering and density of the smectic CHYOs. This is apparent in the hydration profile for the 90:10 system where the water is predominantly co-localized with the ordered ester oxygens of CHYO (Fig. 10a). This ordering may also have implications for the stabilization of class I proteins since the waters are pulled further down into the core making them potentially inaccessible for transmembrane residues close to the PL/core interface.

**Figure 10.**
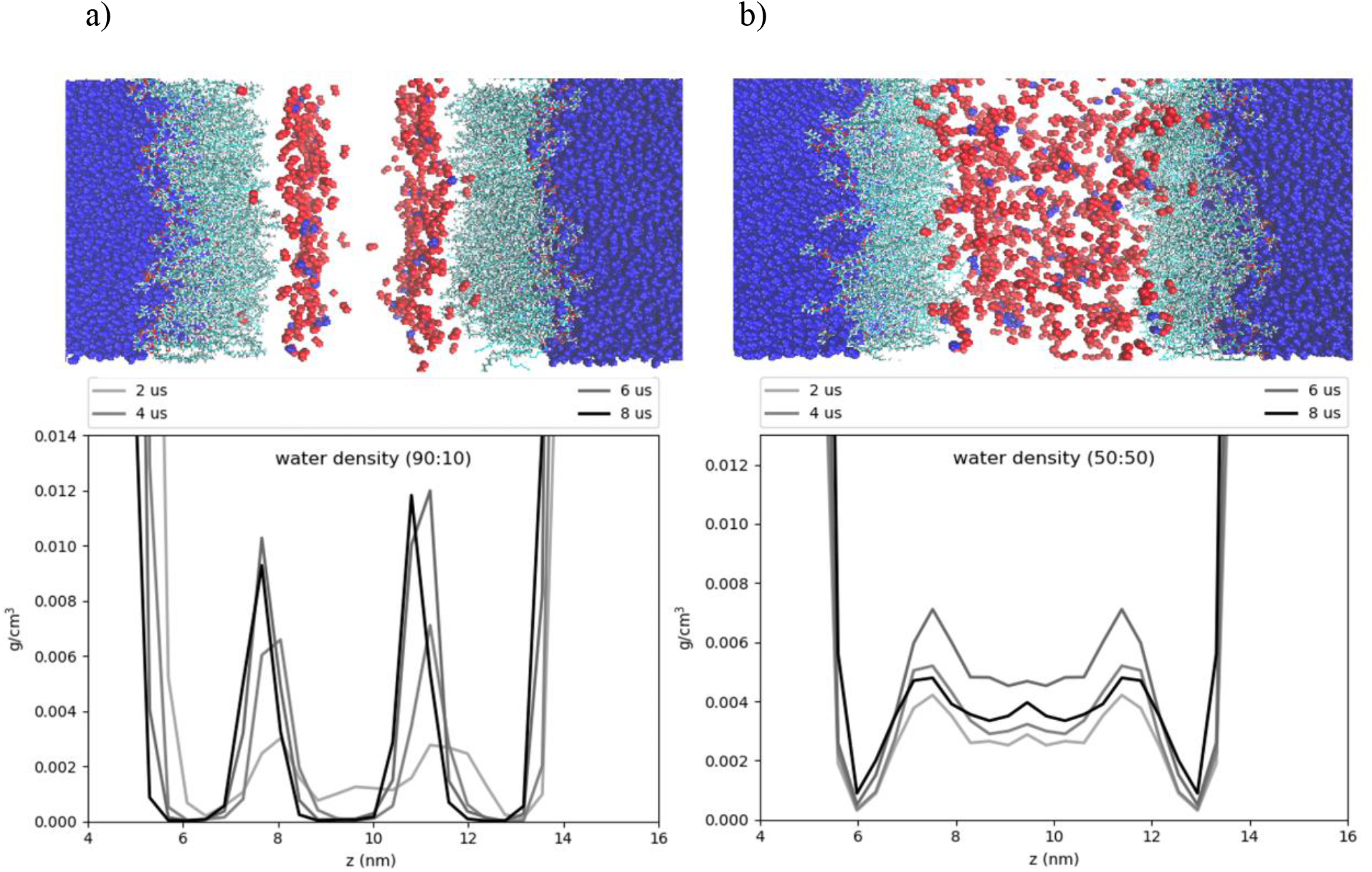
Hydration profiles of the LD systems. The cartoon above the plots shows water (dark blue) phospholipids (cyan), and neutral lipid oxygens (red). The plots display the water density as a function of the z-axis of the systems. The 90:10 ratio system (a) has ordering of the CHYO oxygens, leading to ordering of the hydration profile. (b) The 50:50 ratio system has a more dispersed hydration profile.

To that point, transmembrane motifs are thought to coordinate and make hydrogen bonds with the TG-glycerol oxygens as well.^12^ With CHYO-oleate tails and sidechains existing as the primary chemical component directly below the PL tails for the 90:10 system, potential polar interactions would be significantly reduced. This may contribute to the destabilization of certain proteins as the smectic SE-hydrocarbon tails and chains fill in below the PLs during the phase transition (Fig. S2b).^19^

In summary, our results provide insight into the surface, interfacial, and core physical properties of SE-rich LDs, which vary greatly from those of TG-rich LDs. Namely, with a high ratio of SEs, the LD core undergoes a phase change to a liquid-crystalline smectic phase. This in turn induces ordering in the PL monolayer, reducing the depth that neutral lipids interdigitate and number of SURF-TG molecules, increasing lipid packing and order parameters, decreasing the APL and packing defects, and structuring fewer water molecules in the LD core. CHYO is fundamentally a different molecule from TG and accordingly influences the LD PL monolayer in different ways. The CHYO tails interdigitate just as, if not more readily as TG tails, but they contribute less to packing defects. At lower concentrations they induce less order in the PL tail region than pure TG does (compare 50:50 to pure-TG in Fig. S7), potentially due to CYO’s single oleate tail. However, at sufficiently high concentration the alignment of CHYO into smectic rings translates ordering into the monolayer, significantly reducing the packing defects that play a pivotal role in class II CYTOLD protein association. Collectively, the changes induced by even 50% SE concentration (e.g., changing the percentage and chemical composition of defects) are expected to be sufficient for influencing protein association. Even subtle shifts in the *k*_on_ and *k*_off_ rates of competing proteins may be enough to discriminate association under conditions of competition and crowding. However, the significant changes induced by the amorphous to liquid-crystalline smectic phase transformation under high SE concentrations will have a profound impact on the LD proteome. Understanding the detailed mechanisms of protein association to LDs and how they are impacted by neutral lipid composition will be important topics for future studies.

### Ideas and Speculations

We hypothesize that during times of metabolic stress when cells consume TG and LD cores becomes SE-rich, the metabolic landscape of LDs is transformed via a phase transformation. The resulting liquid-smectic sterol rings under the PL monolayer have substantially altered physical properties that in turn transform the LD proteome. The reduction in surface packing defects will shift the relative stability of class I (ERTOLD) and class II (CYTOLD) proteins and may create a kinetic barrier for protein association that influences the proteome as well. This is supported by class II proteins, such as MLX, that only associate to TG-rich LDs.^5^ The hydrophobic motifs on such proteins will have fewer and smaller packing defects to interact with on surface of SE-rich LDs, and there may be other proteins competing for those defects. Other proteins, such as the class I HSD17B13, that retain association with SE-rich LDs, must rely on interactions with PL head groups or ordered PL tails under more confined conditions. The shift in LD-associated proteins will be governed by a balance of many competing interactions. Hence, even subtle changes could be enough to prioritize one protein over another, especially in the context of enzymes involving slow enzymatic steps following association. Altered off rates could enable selection of certain proteins over others via kinetic proofreading.^46^ Thus, better understanding the association mechanisms of both class I and II proteins to LDs containing different ratios of neutral lipids will be essential to extract trends and insight into the physiochemical control of the LD proteome.

## Methods

### Parameterization of CHYO

We used the CHARMM36 force field for our simulations, due to its history of extensive parametrization and agreement of experimental results for membrane dynamics.^47,48^ The neutral lipids triacylglycerol (TG) and cholesterol oleate (CHYO) were used as model lipids for the LD neutral lipid core. CHARMM36 force field parameters were previously created for TG;^14^ however, the parameters for CHYO did not exist prior to the time of our simulations, meaning parameterization of the molecule was required. Force fields parameters for both cholesterol and oleic acid had previously been established, so we were faced with the task of developing parameters that would unite the two molecules with an ester linkage (Fig. 11a). The initial step was to implement dihedral parameters that are engaged between the two molecule groups as a product of the new ester bond. Quantum mechanical (QM) potential energy surface (PES) scans using the Gaussian 09 software^49^ package were performed on these dihedrals using the MP2/cc-pVDZ basis set^50–52^ as described by previous literature in fitting parameters for CHARMM36 lipid molecules, such as sphingomyelin.^53^ Larger basis sets and treatments of electron correlation have previously been demonstrated to have minimal effect on conformational energies.^53^ Molecular mechanics (MM) PES scans using the GROMACS version 2019.4 simulation engine^54^ were then conducted with the un-parameterized dihedrals, which had their force constants set to 0. These were then subsequently fitted to the QM PES scans using the Monte Carlo Simulated Annealing (MCSA) method, providing force constants for the previously undescribed dihedral parameters.^55^ After fitting, the newly parameterized MM scans agreed with the QM scans, except for a slight underestimation of energy on the peaks (Fig. 11 b,c). Non-bonded parameters were produced from CHARMM General Forcefield Generator (CGenFF), a tool used in CHARMM force fields to fit partial charges among other parameters.^56^ The ester bond and angle types that were needed to connect the cholesterol and oleic acid were already previously defined among the CHARMM36 parameters. To validate the behavior of the newly parameterized cholesteryl oleate, we benchmarked against the only available bulk-phase experimental data: density and phase change properties. At 296K, CHYO has been experimentally demonstrated to have a density of 960 kg/m^3^.^57^ At 1 μs, our simulation of bulk isotropic CHYO under NPT conditions reached a value of 940±9 kg/m^3^. Additionally, pure CHYO, or solutions that contain high ratios of CHYO are also known to undergo a phase transition to a smectic phase at physiological temperatures.^29^ Therefore, another objective was to simulate a phase change, which further validate parameterization efforts. The results for phase changes are discussed in-depth above in the *Results* section above. The topology of CHYO can be found at (https://github.com/jaybraunjr/CHYO-topology).

**Figure 11.**
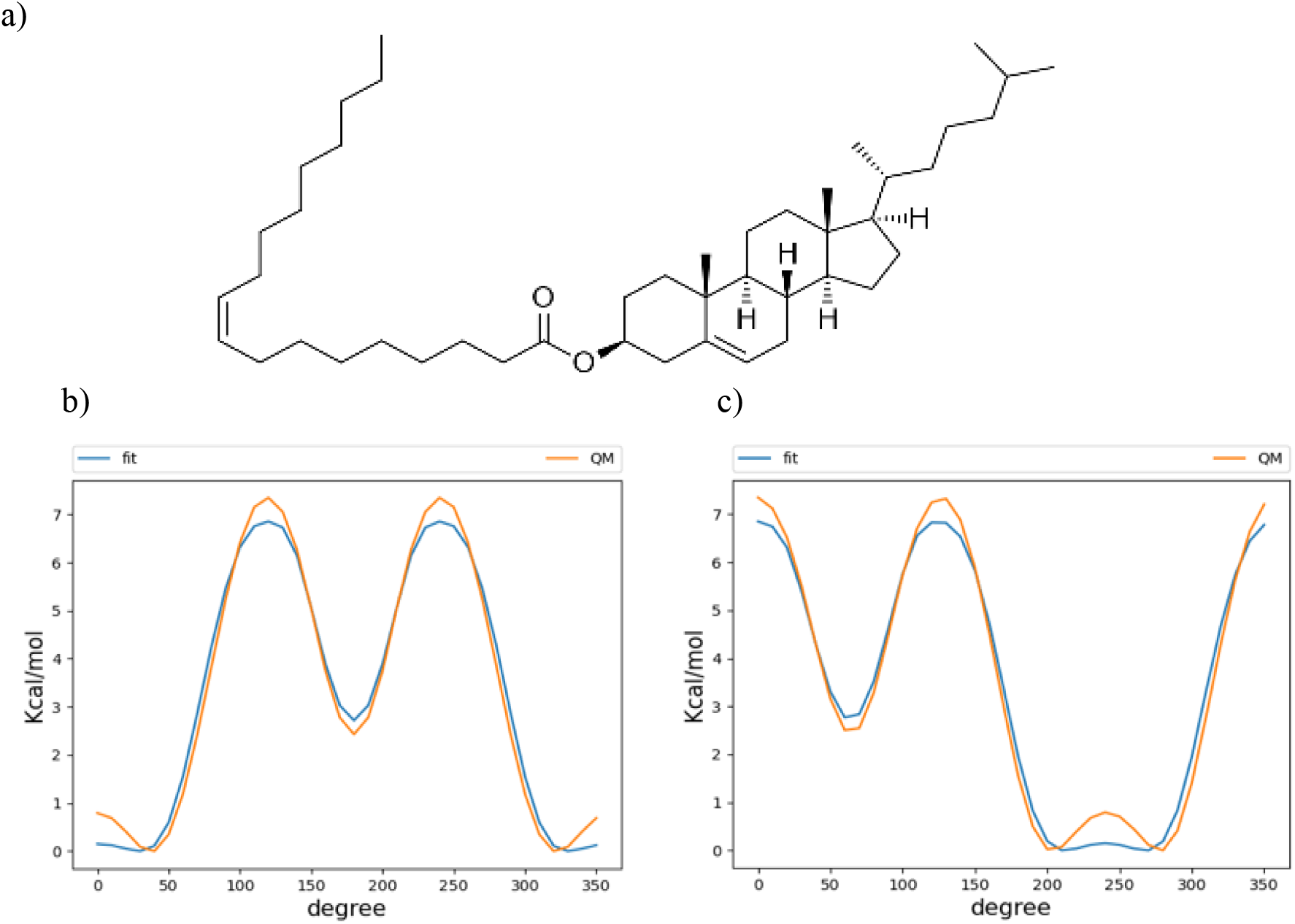
Parameterization of CHYO (a) required fitting of two dihedrals at the ester linkage. The dihedral potential energy scans (b and c) for QM and fitted MM are in agreement, except the MM tends to underestimate energy at the peaks

### System setup

In this work, two separate LD systems were studied: one with a 90:10 CHYO:TG ratio, and another with a 50:50 CHYO:TG ratio. Like previous studies ^14,15^, the LD systems were constructed into trilayer models to allow for atomistic simulations to be conducted with computational efficiency. For a trilayer setup, a bilayer is separated by the top and bottom leaflets, and a neutral lipid core is inserted inside. Water is placed on top and bottom which represents the cytosol. Periodic boundary conditions (PBC) allow the monolayer and core to behave as if the systems were continuous. First, for each LD system, a bilayer was created using the CHARMM-GUI membrane builder ^58^, that contained a heterogeneous population of 88:37:10 amount of the zwitterionic phospholipids: 3-palmitoyl-2-oleoyl-D-glycero-1-phosphatidylcholine (POPC), 2,3-dioleoyl-D-glycero-1-phosphatidylethanolamine (DOPE), and phosphatidylinositol (SAPI), respectively – which is representative of the ER bilayer composition.^59^ The next step was to create a neutral lipid core that matches the *x-y* dimensions of the bilayer, while being thick enough in the *z-*dimension to capture bulk properties and behaviors. Previous simulations^14^ of pure-TG LDs indicated that using a 4 nm thick core led to artefacts such as residual ordering, due to the minimal size in which the TG molecules were allowed to move. Doubling the *z* dimension to create a system with a 8 nm core has been demonstrated to be sufficient to exhibit bulk properties of the neutral lipids.^14^

To create the neutral lipid core, a simulation box with dimensions of approximately 10 × 10 × 4 nm was filled with the neutral lipids CHYO and TG using random placement with the molecular dynamics package Packmol.^60^ 242 neutral lipids were placed in the 90:10 ratio system, and 220 in the 50:50 ratio system. The number of lipids in the respective systems was selected to fill the dimensions of 4 nm thick in the *z* axis, and 10 nm in the *x-y* dimensions to accommodate for the size of the monolayer. The neutral lipid boxes were then equilibrated for 150ns at 310K in NPT and periodic boundary conditions to allow the system to settle. This box was than doubled in the *z*-dimension to create a 8 nm thick core, and it equilibrated again for 100 ns at 310K in NPT with PBC applied. Subsequently, the bilayer created from the CHARMM-GUI output was equilibrated for 10 ns. The bilayer leaflets were separated manually to create monolayers in which the equilibrated neutral lipid box was inserted. An extra 1 nm of space in the *z-*dimensions were allowed between each of the monolayers and neutral lipid box. Neutral lipids that were 2 Å or less from the PLs were removed, and the extra space was reduced with 0.1 ns of NPT simulation. The simulation boxes were solvated with the TIP3 water model^61^, and 0.15 M NaCL solution.

The final systems were then simulated for 8 μs using the GROMACS version 2019.4, with a 2 fs timestep. The temperatures were set to 310K (physiological temperature) using the Nose-Hoover ^62,63^ thermostat and temperature coupling time constant of 1 ps. The particle mesh Ewald (PME) algorithm ^64^ was used to calculate the long-range electrostatic interactions with a cutoff of 1.0 nm. Lennard-Jones pair interactions were cutoff at 12 Å with a force-switching function between 8 and 12Å, and pressure was maintained semi-isotropically using the Parrinello-Rahman barostat.^65^ The pressure was set to 1.0 bar, with a compressibility of 4.5 × 10^−5^ bar^-1^, and a coupling constant of 5.0 ps. The hydrogen bonds were constrained with the LINCS algorithm.^66^ Trajectories were extracted every 100 ps. We analyzed the trajectories using MDAnalysis ^67^, Visual Molecular Dynamics (VMD) ^68^, GROMACS tools, and in-house Python scripting.

### Order parameters

#### Liquid Crystal Ordering

Liquid crystals display properties of both liquids and solids. Essentially, these are substances that aren’t as ordered as a solid yet have some degree of alignment. Therefore, determining the order of the molecules in bulk is useful as a method to quantify respective crystalline-like properties. Liquid crystal meso-phases, such as the nematic and smectic phases, are characterized by molecules aligning in a preferred direction, attaining orientational ordering.^23,69^ The nematic order parameter, which we will refer to as the orientational order parameter (S), quantifies this degree of ordering and alignment in a material.

This is defined as 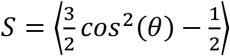 (Equation 1), where *θ* is the angle between the long-axis molecular vector of the molecules and the system director. The system director is the common axis in which the molecules collectively tend to point. The brackets indicate that it is an ensemble average of all the CHYO molecules taken over all the timesteps. In an isotropic liquid, the average of the cosine terms is 0, and therefore the order parameter is equal to 0. On the other hand, when *S* = 1, it indicates a perfect crystalline system.

#### Phospholipid Tail Order Parameters

The tail order parameters use the same formula 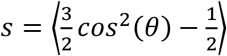, whereas this time *θ* is the angle between the position vector of a carbon atom of an acyl chain bonded to a hydrogen with respect to the system normal (z-axis). A larger value of *S* indicates increased ordering in the membrane. Again, the brackets indicate an ensemble average.^70^

### Classifying surface molecules

The neutral lipids were classified into separate categories based on whether they behave as membrane components or lipids within the LD core. This designation was based on the position of certain atoms with respect to the average *z*-position of the PL tails of the respective monolayer leaflet. If all six oxygens of the TG were above the average *z*-position of the PL tails, then it is considered a SURF-TG. For CHYO, we made this classification (SURF-CHYO) if both ester oxygens were above the average *z*-position of the PL tails. In some cases, only one to several of the designated oxygens would arise past the average *z*-position of the PL tails, however, in these cases most the occurrences were short lived (< 5 ns), and the neutral lipids would return to the bulk of the core. Neutral lipids that exist within the core are referred to as CORE-TG/CHYO.

### Interdigitation

The degree of interdigitation between the neutral lipids and PLs consistent was calculated in a manner consistent with previous studies.^14,15,71^ Interdigitation describes the degree of overlap between neutral lipids and PLs, and it provides us with insight regarding occurrences of membrane-surface properties. This first requires calculating the density profiles of both neutral lipids and PLs with respect to the *z-*axis, which provides us with the overlap parameter:

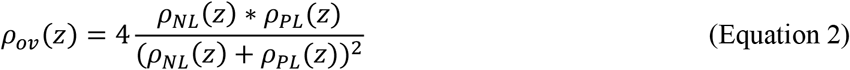

This overlap parameter (*ρ*_*ov*_ (*z*)) ranges from 0 to 1, with 0 meaning that the density profile is equal to zero with no overlap, and 1 meaning there is full overlap (*ρ*_*NL*_(*z*) = *ρ*_*PL*_ (*z*)). To calculate the degree of interdigitation (*λ*_*ov*_), the overlap parameter is then integrated along the z-axis over the whole box:

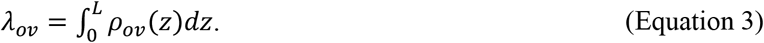

Where *L* is the *z*-dimension of the simulation box, *λ*_*ov*_ is then described in units of nanometers, providing us with an area common to the two density profiles. The interdigitation was calculated for CORE-TG/CHYO, SURF-TG/CHYO, and the two groups combined for total interdigitation. While useful in previous work^14,72^, *λ*_*ov*_ does not effectively differentiate the magnitude of interdigitation for different lipid species in the same system. We therefore also calculated the relative density (*ρ*_*Rel*_) for the PL overlap region, i.e. the region in which the PLs and neutral lipids overlap. This allows for the quantification of different neutral lipid species that interdigitate in the monolayer. The equation for relative density is 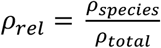, where *ρ* _*total*_ is the total NL + PL density in the overlapping region.

### Packing defect analysis

We employed a Cartesian-based algorithm to calculate the lipid packing defects, adapted from previous work.^14,15,31,73^ For a monolayer leaflet, the lipid atoms whose positions were greater than the threshold (*z*_*thr*_) were projected onto a 1 Åspacing two-dimensional (*x-y*) grid. The threshold (*z*_*thr*_) was set to the average z-position of the phosphorous atoms of the leaflet minus 2 nm, which therefore encompasses most atoms of the respective leaflet. If a grid point overlaps with an atom (center of atom and grid point is less than atom’s radius plus half of the diagonal grid, 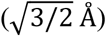, the *z*-position of the atom and the atom type are saved in the grid point. If the grid point overlaps with a polar PL head-group, then it is not considered a defect. If the grid point overlaps with PL-acyl groups or the neutral lipids, then it is considered a defect, with reference to whichever atom has the highest z-position. For each of these defect types, the neighboring elementary defects are clustered into one. If the clustered defect contains N elementary defects, it is considered to have a defect size of N Å^2^. The probability of finding a defect with size N was computed and fit to an exponential decay function: *P*(*N*) = *ce*^−*N*/*π*^, where *c* is the normalization constant and *π* is the packing defect constant. The packing defect constant represents how slowly the decay function falls off. A higher packing defect constant would generally mean that there is a higher probability of finding a larger defect. The script for this packing defect analysis can be found at (https://github.com/ksy141/SMDAnalysis).

## Acknowledgements

The authors thank Dr. Siyoung Kim, Professor Rich Pastor and Professor Jeffrey Klauda for helpful discussions. We gratefully acknowledge support from the National Institute of General Medicine of the National Institutes of Health under award number R35GM143117 and computational support from the Extreme Science and Engineering Discovery Environment supported by the National Science Foundation (Grant No. ACI-1548562) under allocation MCB200018 as well as the Center for High Performance Computing at the University of Utah.

## Notes

### Competing Interest Statement

The authors have declared no competing interest.

