## Supplementary material for "Capturing the liquid-crystalline phase transformation: Implications for protein targeting to sterol ester-rich lipid droplets": SI

a) b)

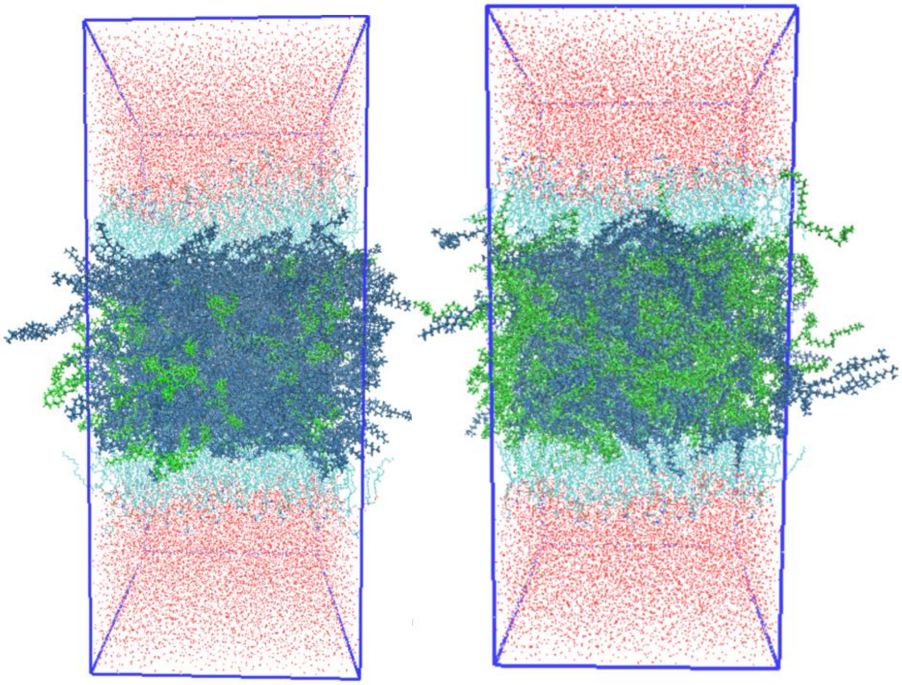

Figure S1. (a) 90:10 CHYO:TG system, and (b) 50:50 CHYO:TG system during the 1st μs of simulation, with neutral lipids arranged randomly. CHYO is blue, TG is green. Trilayer systems are used to represent LD's, see *Methods* section for more details.

b) b)

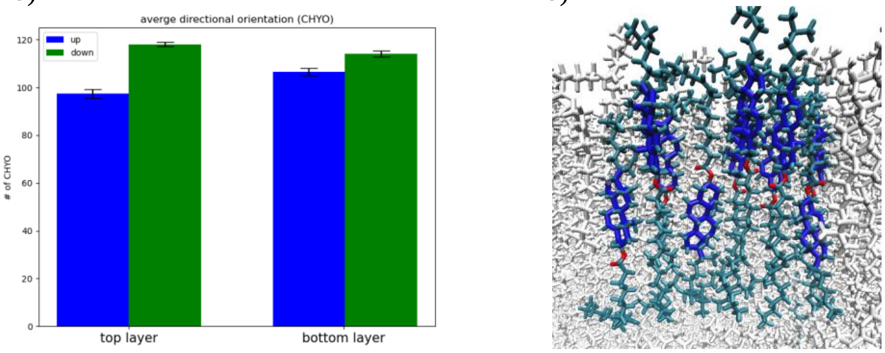


Figure S2. Calculation of the orientation of CHYO molecules per smectic layer (a), and a screenshot of CHYO in one of the layers displaying the alternating orientation (b).

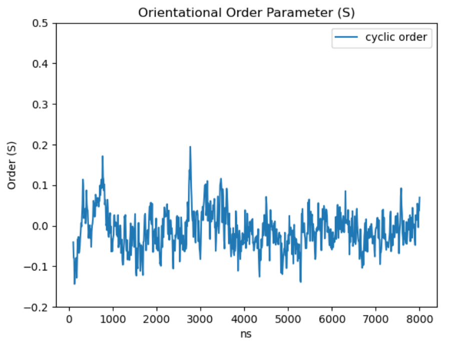


Figure S3. The orientational order parameter (S) for the 50:50 CHYO:TG LD averages 0, indicating no ordering has occurred.

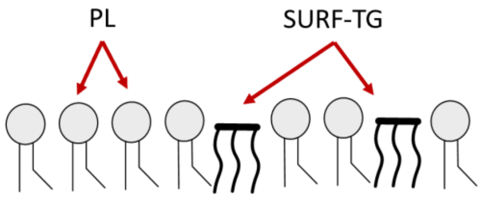


Figure S4. Representation of SURF-TGs in the PL monolayer. The SURF-TGs behave as membrane components

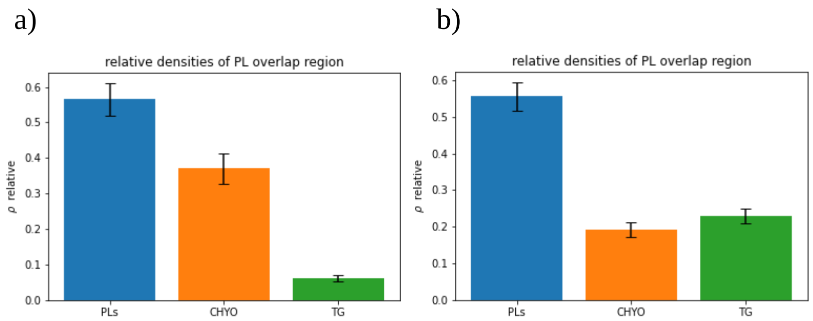


Figure S5. The interdigitation of the neutral lipids and phospholipids. (a) $\rho_{rel}$ for the 90:10 system and (b) $\rho_{rel}$ for the 50:50 system.

Table S1. The average interdigitation depth, $\lambda_{OV}$ , of neutral lipids. The calculations were taken after 4 μs to allow for equilibration.

| **LD type** | **90:10 CHYO:TG LD** | **50:50 CHYO:TG LD** | **Pure TG LD** |
|---|---|---|---|
| SURF-TG | 0.130±0.014 | 0.605±0.039 | ~1.27 |
| CORE-TG | 1.188±0.155 | 1.657±0.139 | ~1.83 |
| TG total | 1.294±0.157 | 1.998±0.0.127 | ~2.42 |
| CHYO | 1.278±0.097 | 1.678±0.129 | |

***Standard errors SURF:*** *90:10 TG: 0.001, 50:50 TG: 0.001*
***Standard errors CORE:*** *90:10 TG: 0.008, 50:50 TG: 0.007*
***Standard errors total:*** *90:10 CHYO: 0.005, 50:50 CHYO: 0.008, 90:10 TG: 0.008, 50:50 TG: 0.007*

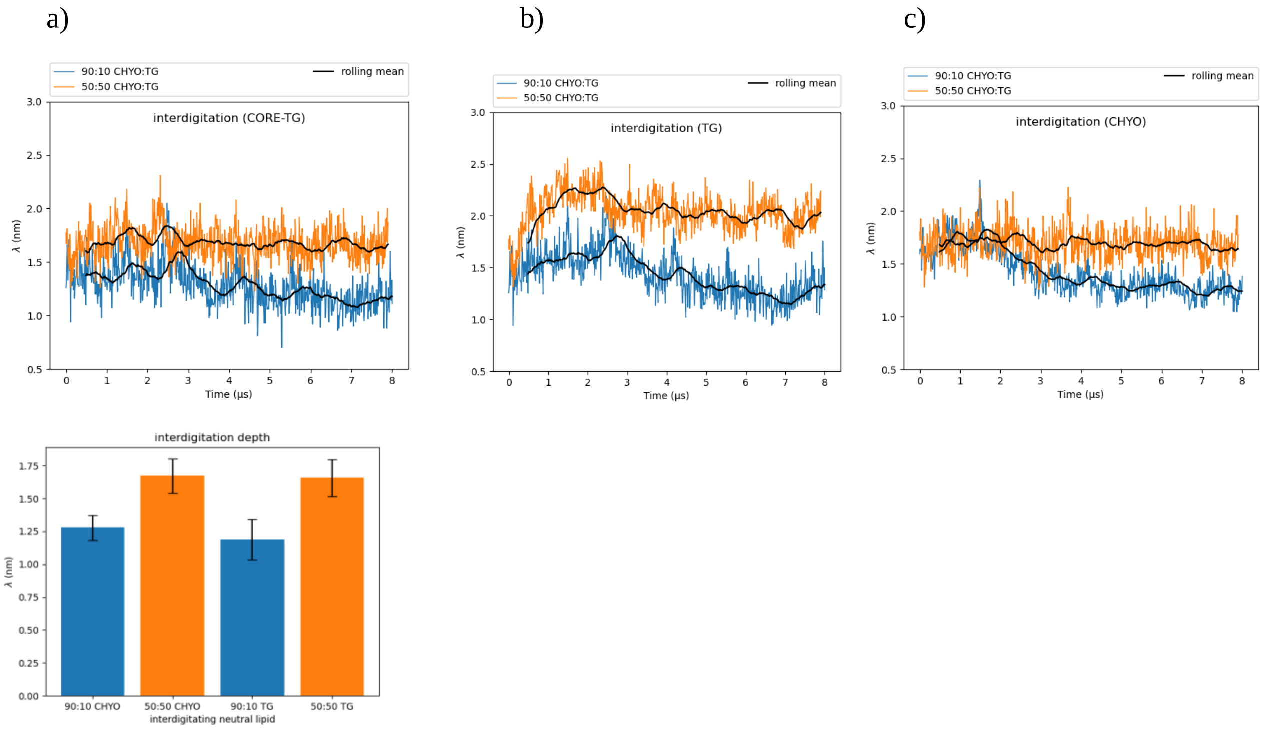


Figure S6. (a) Interdigitation CORE-TG, (b) total-TG, and (c) total-CHYO. (d) Average total interdigitation depth after initial equilibration of 3 μs.

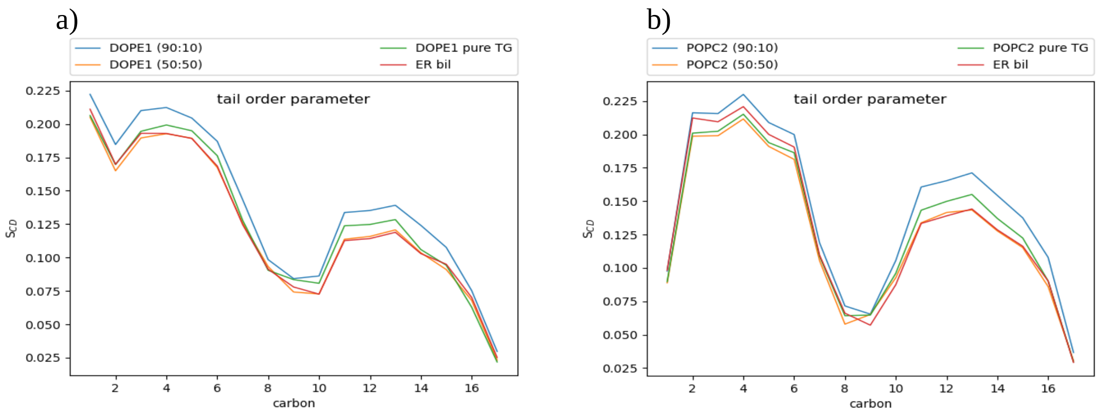


Figure S7. PL tail order parameters for (a) DOPE and (b) POPC. In these plots, we included the order parameters of an ER bilayer (red) and a pure-TG LD (green) for comparison purposes.
